# Mathematical Characterization of Private and Public Immune Repertoire Sequences

**DOI:** 10.1101/2022.05.17.492232

**Authors:** Lucas Böttcher, Sascha Wald, Tom Chou

**Author notes:** Submitted to the editors on May 9, 2022. **Funding:** LB received funding from the Swiss National Fund (P2EZP2 191888). TC acknowledges funding from the NIH through grant R01HL146552 and the NSF through grant DMS-1814364.

## Abstract

Diverse T and B cell repertoires play an important role in mounting effective immune responses against a wide range of pathogens and malignant cells. The number of unique T and B cell clones is characterized by T and B cell receptors (TCRs and BCRs), respectively. Although receptor sequences are generated probabilistically by recombination processes, clinical studies found a high degree of sharing of TCRs and BCRs among different individuals. In this work, we formulate a mathematical and statistical framework to quantify receptor distributions. We define information-theoretic metrics for comparing the frequency of sampled sequences observed across different individuals. Using synthetic and empirical TCR amino acid sequence data, we perform simulations to compare theoretical predictions of this clonal commonality across individuals with corresponding observations. Thus, we quantify the concept of “publicness” or “privateness” of T cell and B cell clones. Our methods can also be used to study the effect of different sampling protocols on the expected commonality of clones and on the confidence levels of this overlap. We also quantify the information loss associated with grouping together certain receptor sequences, as is done in spectratyping.

## 1. Introduction

A major component of the adaptive immune system in most jawed vertebrates is the repertoire of B and T lymphocytes. A diverse immune repertoire allows the adaptive immune system to recognize a wide range of pathogens [44]. B and T cells develop from common lymphoid progenitors (CLPs) that originate from hematopoietic stem cells (HSCs) in the bone marrow. B cells mature in the bone marrow and spleen while developing T cells migrate to the thymus where they undergo their maturation process. After encountering an antigen, naive B cells may get activated and differentiate into antibody-producing plasma cells, which are essential for humoral (or antibody-mediated) immunity. In recognizing and eliminating infected and malignant cells, T cells contribute to cell-mediated immunity of adaptive immune response.

T-cell receptors bind to antigen peptides (or epitopes) that are presented by major histocompatibility complex (MHC) molecules on the surface of antigen-presenting cells (APCs). T cells that each carry a type of TCR mature in the thymus and undergo V(D)J recombination, where variable (V), diversity (D), and joining (J) gene segments are randomly recombined [3, 41]. The receptors are heterodimeric molecules and mainly consist of an *α* and a *β* chain while only a minority, about 1–10% [20], of TCRs consists of a *δ* and a *γ* chain. The TCR *α* and *γ* chains are made up of VJ and constant (C) regions. Additional D regions are present in *β* and *γ* chains. During the recombination process, V(D)J segments of each chain are recombined randomly with additional insertions and deletions. After recombination, only about 5% or even less [45] of all generated TCR sequences are selected based on their ability to bind to certain MHC molecules (“positive selection”) and to not trigger autoimmune responses (“negative selection”). These naive T cells are then exported from the thymus into peripheral tissue where they may interact with epitopes that are presented by APCs. The selection process as well as subsequent interactions are specific to an individual.

The most variable parts of TCR sequences are the complementary determining regions (CDRs) 1, 2, and 3, located within the V region, among which the CDR3*β* is the most diverse [2]. Therefore, the number of distinct receptor sequences, the richness *R*, of TCR repertoires is typically characterized in terms of the richness of CDR3*β* sequences. Only about 1% of T cells express two different TCR*β* chains [14, 33, 37], whereas the proportion of T cells that express two different TCR*α* chains may be as high as 30% [36, 37].

B cells can also respond to different antigens via different B cell receptors (BCRs) that are comprised of heavy and light chains. As with TCRs, the mechanism underlying the generation of a diverse pool of BCRs is VDJ recombination in heavy chains and VJ recombination in light chains. Positive and negative selection processes sort out about 90% of all BCRs that react too weakly or strongly with certain molecules [42]. As a result of the various recombination and joining processes and gene insertions and deletions, the theoretically possible richness of BCR and TCR receptors is about 10^14^ − 10^15^ [46, 32]. However, the actual number of unique receptors (or immunotypes/clonotypes) realized in organisms is much smaller. The richness of TCR sequences was estimated to be about 10^6^ for mice [6] and about 10^8^ for humans [40]. B cell richness for humans is estimated to be 10^8^ − 10^9^ [16]. Estimating the true richness of BCR and TCR pools in an organism is challenging since the majority of such analyses are based on small blood samples, leading to problems similar to the “unseen species” problem in ecology [31].

Each pool of BCR and TCR sequences realized in one organism *i* can be seen as a subset 𝒰_*i*_ of the set of all possible species-specific sequences 𝒮. Sequences that occur in at least two different organisms *i* and *j* (*i*.*e*., sequences that are elements of 𝒰_*i*_∩𝒰_*j*_) are commonly referred to as “public” sequences [31] while “private” sequences occur only in one of the individuals tested. The existence of public TCR*β* sequences has been established in several previous works [34, 35, 38, 40]. More recently, a high degree of shared sequences has been also observed in human BCR repertoires [5, 39].

The notions of public and private clonotypes are only loosely defined. Some references use the term “public sequence” to refer to those sequences that “are *often* shared between individuals” [38] or “shared across individuals” [25]. In this work, we provide more quantitative definitions of “public” and “private” clones for general clone size-distributions and *M* individuals.

In the next section, we first give an overview of the mathematical concepts that are relevant to characterize TCR and BCR distributions. We then formulate a statistical model of receptor distributions in Sec. 3. In Secs. 3.1 and 3.2, we derive quantities associated with receptor distributions in single organisms and across individuals, respectively. We will primarily focus on the overlap of repertoires across individuals and on the corresponding confidence intervals that can be used to characterize “public” and “private” sequences of immune repertoires. Formulae we derived are listed in Table 1. In Sec. 4, we use synthetic and empirical TCR amino acid sequence data and perform simulations to compare theoretical predictions of repertoire overlaps between different individuals with corresponding observations. Our source codes are publicly available at [1].

**TABLE 1:** Table of mathematical results. We list our main mathematical derivations and expressions for richness and overlap, unsampled and sampled, in both the ***n***-representation and the ***p***-representation. The component probabilities 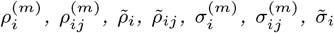, and 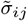 are given in Eqs. (3.3), (3.5), (3.10), (3.13), (3.22), (3.23), (3.24), and (3.25), respectively. The component probabilities 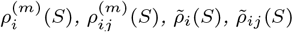 associated with sampled quantities in the ***p***-representation are given by Eqs. (3.35), (3.36), (3.37), and (3.38), respectively.

| Metric | Description | n-representation | p-representation |
| --- | --- | --- | --- |
| $\mathbb{E}[R]$ | individual richness | $\sum_{k=1}^N \sum_{i=1}^{\Omega} \mathbb{1}(n_i, k)$ | $\sum_{i=1}^{\Omega} \rho_i$ |
| $\mathbb{E}[R^2]$ | 2<sup>nd</sup> moment of richness | $\left[ \sum_{k=1}^N \sum_{i=1}^{\Omega} \mathbb{1}(n_i, k) \right]^2$ | $\sum_{i=1}^{\Omega} \rho_i + \sum_{i \neq j}^{\Omega} \rho_{ij}$ |
| $\mathbb{E}[R_s]$ | mean sampled richness | $\sum_{i=1}^{\Omega} \sigma_i = \Omega - \frac{1}{\binom{N}{S}} \sum_{i=1}^{\Omega} \binom{N-n_i}{S}$ | $\sum_{i=1}^{\Omega} \rho_i$ |
| $\mathbb{E}[R_s^2]$ | 2<sup>nd</sup> moment, sampled richness | $\sum_{i=1}^{\Omega} \sigma_i + \sum_{i \neq j}^{\Omega} \sigma_{ij}$ | $\sum_{i=1}^{\Omega} \rho_i(S) + \sum_{i \neq j}^{\Omega} \rho_{ij}(S)$ |
| $\mathbb{E}[R^{(M)}]$ | mean group richness | $\sum_{k \geq 1} \sum_{i=1}^{\Omega} \mathbb{1}(\sum_{m=1}^M n_i^{(m)}, k)$ | $\sum_{i=1}^{\Omega} \tilde{\rho}_i$ |
| $\mathbb{E}[(R^{(M)})^2]$ | 2<sup>nd</sup> moment, grp richness | $\left[ \sum_{k \geq 1} \sum_{i=1}^{\Omega} \mathbb{1}(\sum_{m=1}^M n_i^{(m)}, k) \right]^2$ | $\sum_{i=1}^{\Omega} \tilde{\rho}_i + \sum_{i \neq j}^{\Omega} \tilde{\rho}_{ij}$ |
| $\text{Var}[R^{(M)}]$ | variance of grp richness | 0 | $\sum_{i \neq j}^{\Omega} \tilde{\rho}_{ij} + \left( 1 - \sum_{i=1}^{\Omega} \tilde{\rho}_i \right) \sum_{i=1}^{\Omega} \tilde{\rho}_i$ |
| $\mathbb{E}[R_s^{(M)}]$ | mean sampled grp richness | $\sum_{i=1}^{\Omega} \tilde{\sigma}_i$ | $\sum_{i=1}^{\Omega} \tilde{\rho}_i(S)$ |
| $\mathbb{E}[(R_s^{(M)})^2]$ | 2<sup>nd</sup> moment, sampled grp richness | $\sum_{i=1}^{\Omega} \tilde{\sigma}_i + \sum_{i \neq j}^{\Omega} \tilde{\sigma}_{ij}$ | $\sum_{i=1}^{\Omega} \tilde{\rho}_i(S) + \sum_{i \neq j}^{\Omega} \tilde{\rho}_{ij}(S)$ |
| $\mathbb{E}[K^{(M)}]$ | expected $M$-overlap | $\sum_{i=1}^{\Omega} \prod_{m=1}^M \sum_{k^{(m)} \geq 1} \mathbb{1}(n_i^{(m)}, k^{(m)})$ | $\sum_{i=1}^{\Omega} \prod_{m=1}^M \rho_i^{(m)}$ |
| $\mathbb{E}[(K^{(M)})^2]$ | 2<sup>nd</sup> moment, $M$-overlap | $\left[ \sum_{i=1}^{\Omega} \prod_{m=1}^M \sum_{k^{(m)} \geq 1} \mathbb{1}(n_i^{(m)}, k^{(m)}) \right]^2$ | $\sum_{i=1}^{\Omega} \prod_{m=1}^M \rho_i^{(m)} + \sum_{i \neq j}^{\Omega} \prod_{m=1}^M \rho_{ij}^{(m)}$ |
| $\mathbb{E}[K_s^{(M)}]$ | sampled $M$-overlap | $\sum_{i=1}^{\Omega} \prod_{m=1}^M \sigma_i^{(m)}$ | $\sum_{i=1}^{\Omega} \prod_{m=1}^M \rho_i^{(m)}(S)$ |
| $\mathbb{E}[(K_s^{(M)})^2]$ | 2<sup>nd</sup> moment, sampled $M$-overlap | $\sum_{i=1}^{\Omega} \prod_{m=1}^M \sigma_i^{(m)}(S) + \sum_{i \neq j}^{\Omega} \prod_{m=1}^M \sigma_{ij}^{(m)}(S)$ | $\sum_{i=1}^{\Omega} \prod_{m=1}^M \rho_i^{(m)}(S) + \sum_{i \neq j}^{\Omega} \prod_{m=1}^M \rho_{ij}^{(m)}(S)$ |

## 2. Mathematical concepts

Although receptor sequences and cell counts are discrete quantities, using continuous functions to describe their distribution may facilitate the mathematical analysis of the quantities that we derive in the subsequent sections. We therefore briefly describe some elementary concepts associated with the discretization of continuous distributions.

Let *p*(*x*) be the *probability density* associated with the distribution of traits, as depicted in Fig. 1(a). The probability that a certain trait occurs in [*x, x* + d*x*) is *p*(*x*) d*x*. The corresponding discretized distribution elements are

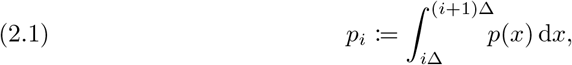

where Δ is the discretization step size of the support of *p*(*x*). If we discretize the values of probabilities, the number of clones with a certain relative frequency *p*_*i*_ is given by the *clone count*

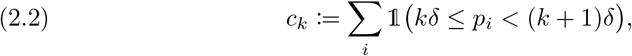

where the indicator function 𝟙 = 1 if its argument is satisfied and zero otherwise. For *x, y* ∈ ℤ_≥0_, we also employ the notation 𝟙(*x, y*) = 1 if *x* = *y* and 𝟙(*x, y*) = 0 if *x* ≠ *y*. As shown in Fig. 1(b), the parameter *δ* defines an **interval** of frequency values and modulates the clone-count binning. Figures 1(b,c) show how *p*_*i*_ and *c*_*k*_ are constructed from a continuous distribution *p*(*x*).

**FIG. 1.**
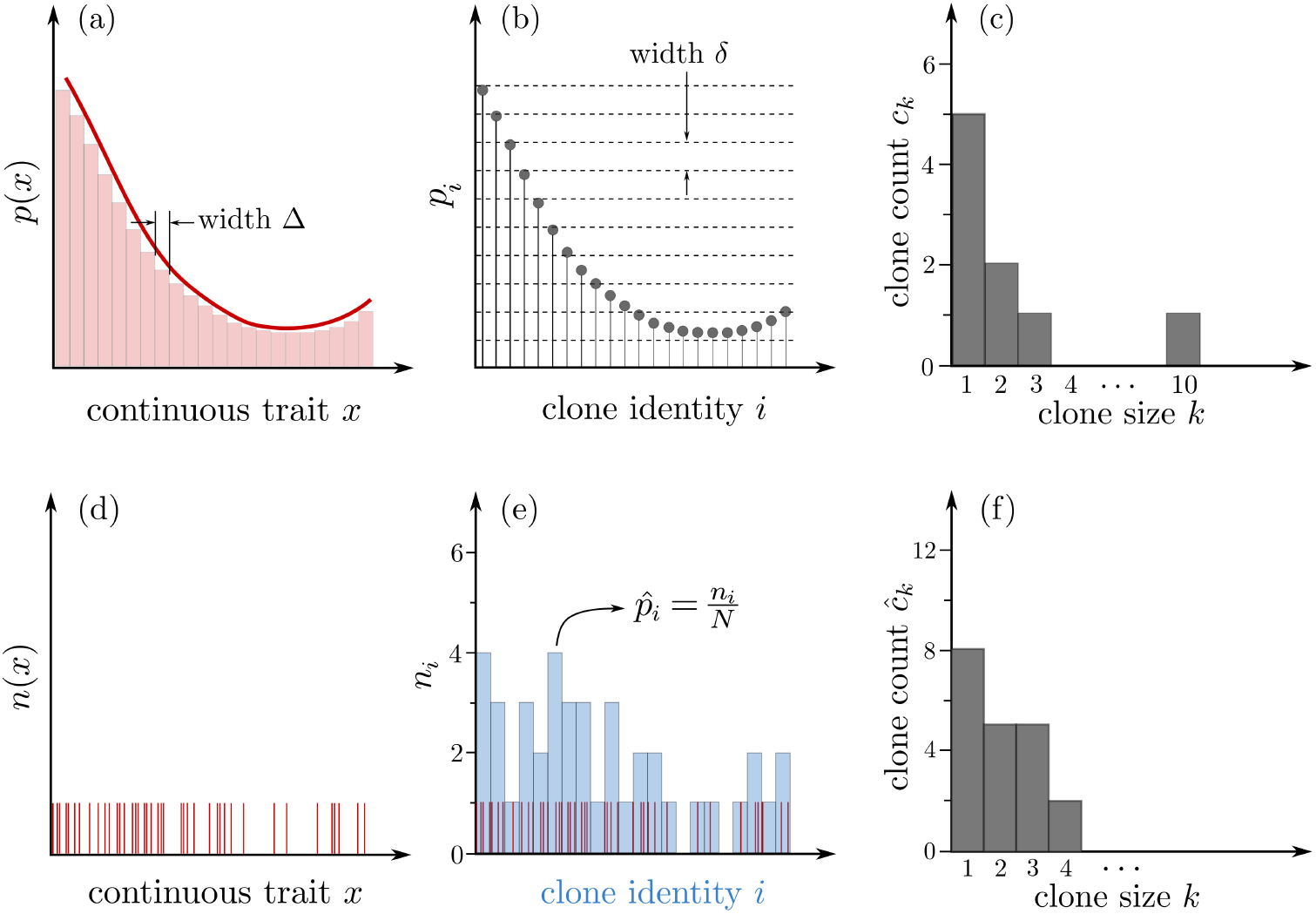
Sampling from a continuous distribution, described in terms of an underlying probability density p(x) and number density n(x). The probability density p(x) (solid red line) and the Riemann sum approximation to the probability (red bars of width *Δ*) are shown in panel (a). The probability that a trait in the interval *[x, x + dx)* arises is p(x)dx. As shown in (b), this distribution can be discretized directly by the intervals *[iΔ, (i + 1)Δ)* (red bars) defining discrete traits and their associated probabilities p_i_ (see Eq. (2.1)). The probabilities p_i_ can be transformed into clone counts c_k_ (the number of identities i that are represented by k individuals) using Eq. (2.2), and are shown in (c). A finite sample of a population described by p(x) yields the binary outcome shown in (d). In this example, the total number of samples is N = 41 and and since the trait space x is continuous, the probability that the exact same trait arises in more than one sample is almost surely zero. Light blue bars in panel (e) represent number counts n_i_ binned according to *Δ*. The probabilities 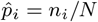 provide an approximation of p_i_. Clone counts for the empirical 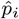 are calculated according to Eq. (2.3) and shown in (f).

If *p*(*x*) is not available from data or a model, an alternative representative starts with the number density *n*(*x*), which can be estimated by sampling a process which follows *p*(*x*). The probability that a continuous trait *x* is drawn twice from a continuous distribution *p*(*x*) is almost surely zero. Hence, the corresponding number counts *n*(*x*) are either 1 if *X* ∈ [*x, x*+d*x*) (*i*.*e*., if trait *X* is sampled) or 0 otherwise, as shown in Figs. 1(d,e). We say that *X* is of *clonotype i* if *X* ∈ [*i*Δ, (*i* + 1)Δ) (1 ≤ *i* ≤ Ω) and we use *n*_*i*_ to denote the number of cells of clonotype *i*. Then, if Ω denotes the theoretical maximum number of different clonotypes, the total T cell (or B cell) population is 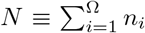. The relative empirical abundance of clonotype *i* is thus 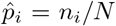 (see Fig. 1d), satisfying the normalization condition 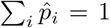. The corresponding empirical clone count derived from the number representation *n*_*i*_ is defined as

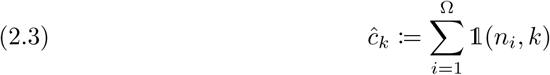

and shown in Fig. 1(f). Besides the simple discrete estimate 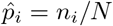, one can also reconstruct *p*(*x*) from **n** = {*n*_*i*_} using methods such as kernel density estimation.

## 3. Whole organism statistical model

Using the mathematical quantities defined above, we develop a simple statistical model for BCR and TCR sequences distributed among individuals. Although our statistical model is applicable to both BCR and TCR sequences, we will primarily focus on the characterization of TCRs for simplicity. For example, B cells undergo an additional process of somatic hypermutation and class switching leading to a more dynamic evolution of the more diverse B cell repertoire [18]. By focusing on naive T cells, we can assume their populations are generated by the thymus via a single, simple effective process.

Assume a common universal recombination process (see Fig. 2) in T cell development that generates a cell with TCR of type 1 ≤ *i* ≤ Ω with probability 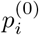. The theoretical number of possible sequences before thymic selection is very large, Ω ∼ 10^16^. Although each new T cell produced carries TCR *i* with probability 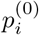, many of these probabilities are effectively zero given both selection (that eliminates ∼ 98% of them) and the finite number of T cells produced over a lifetime [41, 45, 32].

**FIG. 2.**
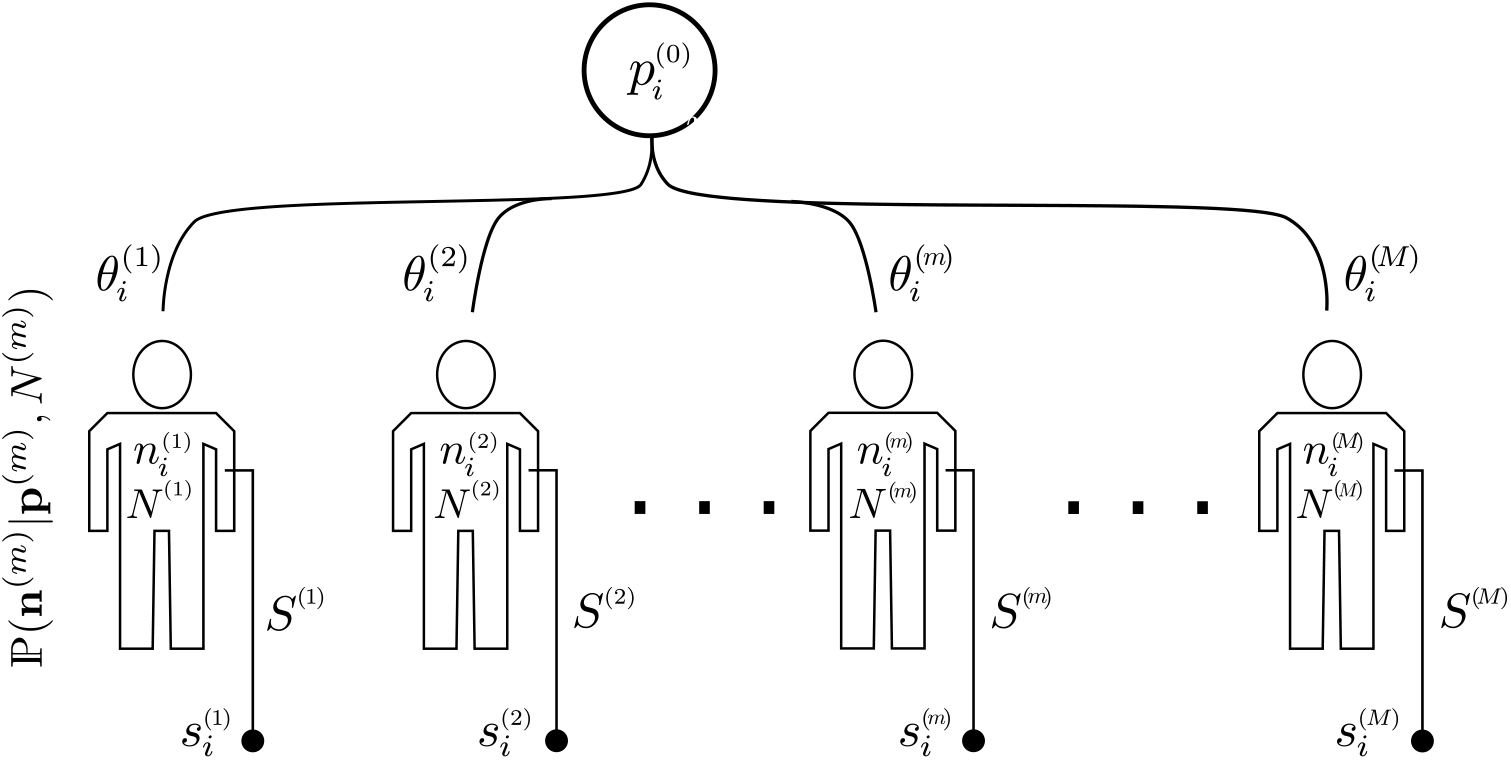
Schematic of sampling of multiple species from multiple individuals. A central process produces (through V(D)J recombination) TCRs. Individuals select for certain TCRs resulting in population 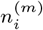 of T cells with receptor i in individual m, for a total T cell count 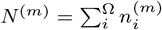. The selection of TCR i by individual m (in their individual thymuses) is defined by the parameter 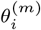 which gives an effective probability 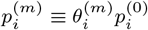. A sample with cell numbers S^(m)^ ≪ N ^(m)^ is drawn from individual m and sequenced to determine 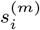, the number of cells of type i in the subsample drawn from individual m.

Besides thymic selection and subsequent death, activation, and proliferation occur differently across individuals 1 ≤ *m* ≤ *M*, described by model parameters 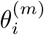. Such a model translates the fundamental underlying recombination process into a population of 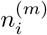 T cells with TCR *i* and total population 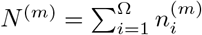 in individual *m*. The connection between 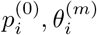 and 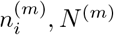 might be described by population dynamics models, deterministic or stochastic, such as those presented in [15].

At any specific time, individual *m* will have a cell population configuration 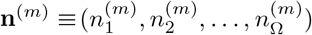 with probability ℙ(**n**^(*m*)^|***θ***^(*m*)^, *N* ^(*m*)^). Each individual can be thought of as a biased sample from all cells produced via the universal probabilities 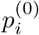. We can define individual probabilities 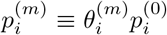 and model the probability of a T cell population **n**^(*m*)^ in individual *m* by a multinomial distribution over individual probabilities 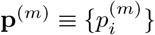:

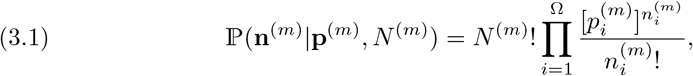

where 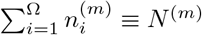 and 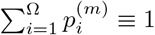.

Thus, each individual can be thought of as a “sample” of the “universal” thymus. Neglecting genetic relationships amongst individuals, we can assume them to be independent with individual probabilities 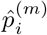, the effective probability at which a cell of type *i* is generated in individual *m*. An estimate of this individual probability is 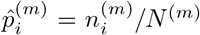. A dynamical model for 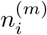 would explicitly involve activation, proliferation, and death rates of different classes of T cells. A set of probabilities 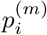 to describe probabilities of immigration of a type *i* cell is chosen out of convenience. This representation allows us to more simply analytically express the probabilities of any configuration **n**^(*m*)^. A detailed mechanistic model may be used to relate its parameters to 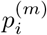.

### 3.1. Single individual quantities

First we focus on quantities intrinsic to a single individual organism; thus, we can suppress the “*m* = 1” label. Within an individual, we can use clone counts to define metrics such as the richness

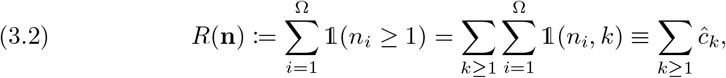

where 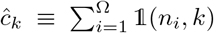 is defined in Eq. (2.3) (the number of clones that are of population *k*). Other diversity/entropy metrics such as Simpson’s indices, Gini indices, etc. [44] can also be straightforwardly defined. Given the clone populations **n**, the individual richness can be found by direct enumeration of Eq. (3.2).

We can also express the richness in terms of the underlying probabilities **p** associated with the individual by first finding the probability that a type-*i* cell appears at all among the *N* cells within said individual

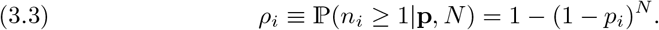

This probability is associated with a binary outcome, either appearing or not appearing. Higher order probabilities like *ρ*_*ij*_ (both *i*- and *j*-type cells appearing in a specific individual) can be computed using the marginalized probability

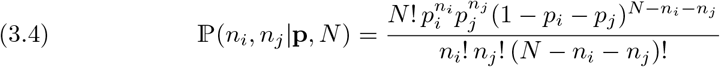

to construct

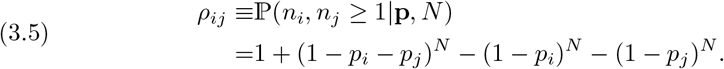

Higher moments can straightforwardly computed using quantities such as

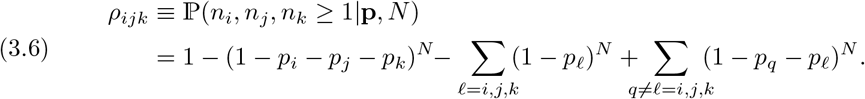

These expressions arise when we compute the moments of *R* [defined by Eq. (3.2)] in terms of the probabilities **p**. Using the single-individual probability ℙ(**n**|**p**, *N*) (Eq. (3.1)) allows us to find moments of the richness in a single individual in terms of **p**, with the first two given by

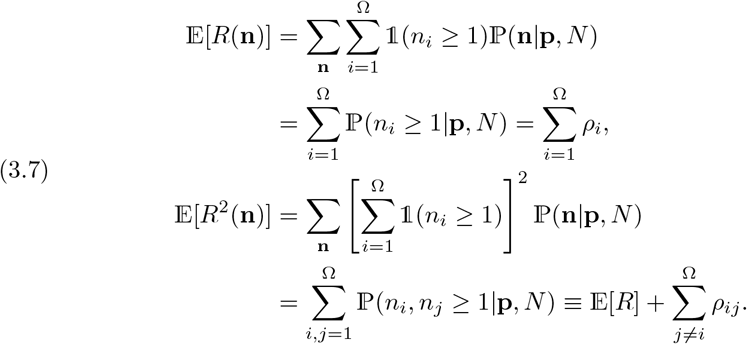

### 3.2. Multi-individual quantities

Here, we consider the distribution **n**^(*m*)^ across different individuals and construct quantities describing group properties. For example, the combined richness of all TCR clones of *M* individuals is defined as

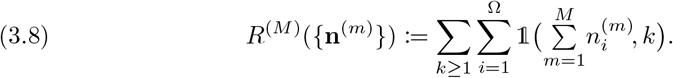

To express the expected multi-individual richness in terms of the underlying individual probabilities **p**^(*m*)^, we weight Eq. (3.8) over the *M*-individual probability

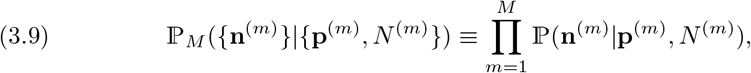

and sum over all allowable **n**^(*m*)^. For computing the first two moments of the total-population richness in terms of **p**^(*m*)^, we will make use of the marginalized probability 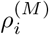 that clone *i* appears in at least one of the *M* individuals

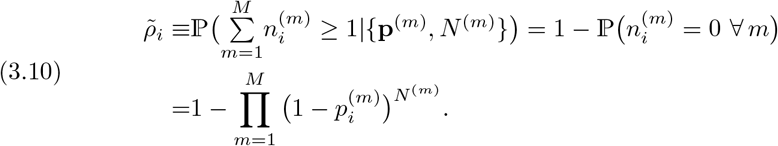

We will also need the joint probability that clones *i* and *j* both appear in at least one of the *M* individuals 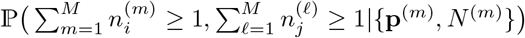, which we can decompose as

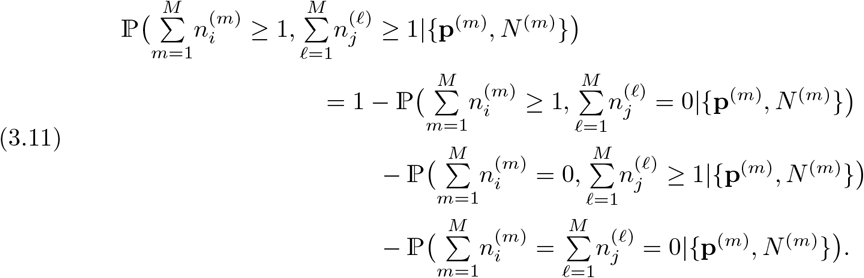

Upon using Eqs. (3.1) and (3.4), we find

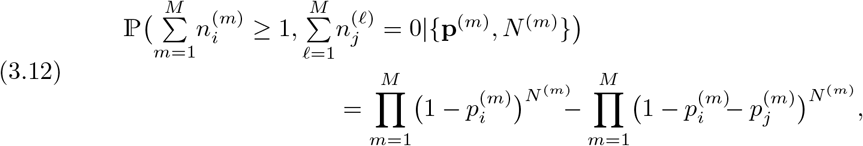

allowing us to rewrite Eq. (3.11) as

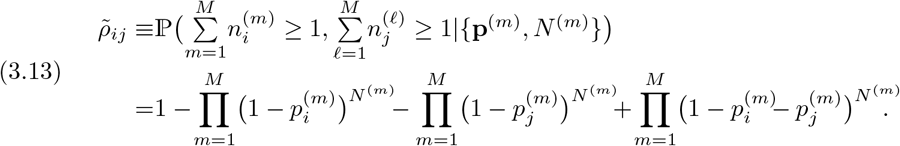

Note that 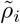 and 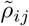 are different from 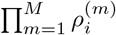 and 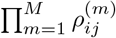,

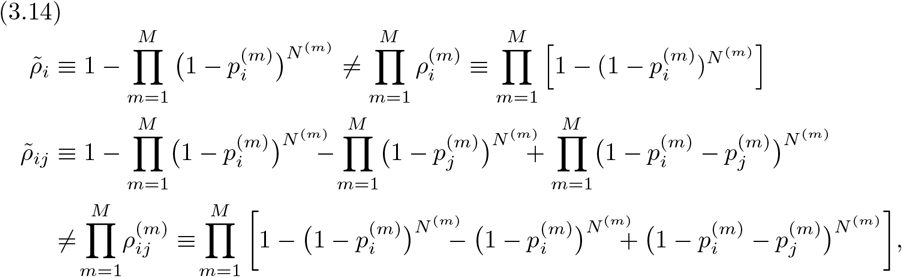

with, *e*.*g*., 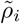 describing the probability that a type *i* cell occurs at all in the total population, while 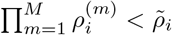 describes the probability that a type *i* cell appears in each of the *M* individuals.

Using the above definitions, we can express the mean total-population richness as

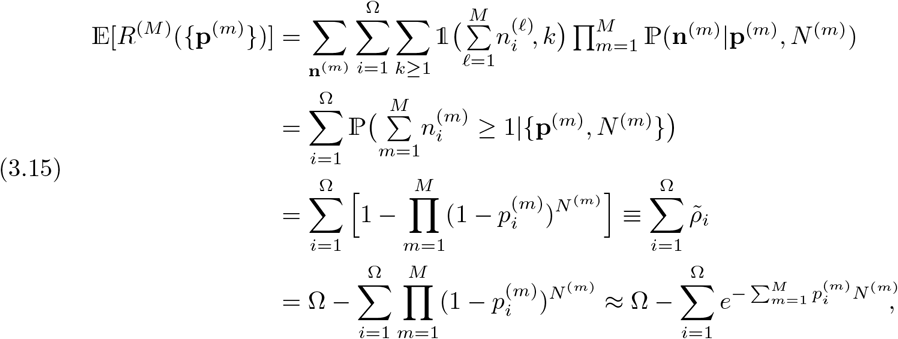

where the last approximation holds for 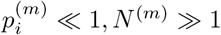. The second moment of the total *M*-population richness can also be found in terms of 𝔼[*R*^(*M*)^] and Eq. (3.13),

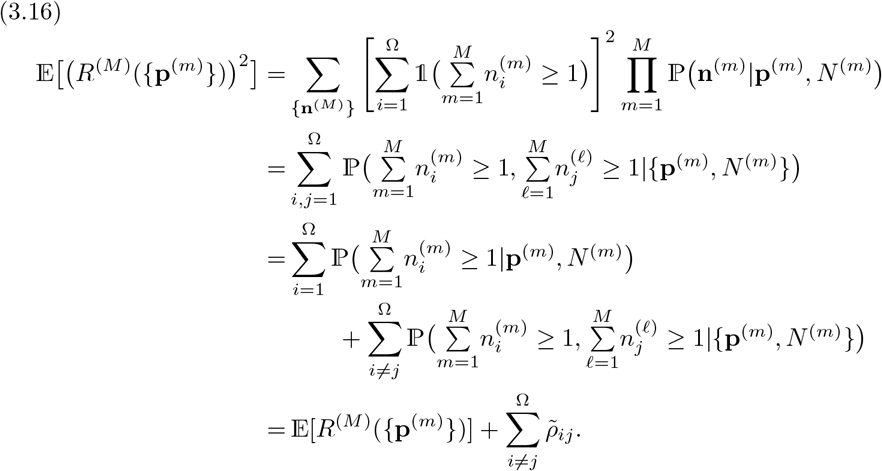

Given **n**^(*m*)^ of all individuals, we can also easily define the number of distinct TCR clones that appear in all of *M* randomly selected individuals, the “overlap” or “*M*-publicness”

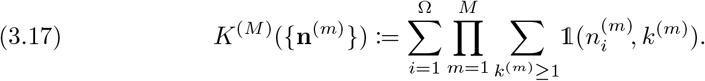

Figure 3 provides a simple example of three individuals each with a contiguous distribution of cell numbers 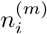 that overlap.

**FIG. 3.**
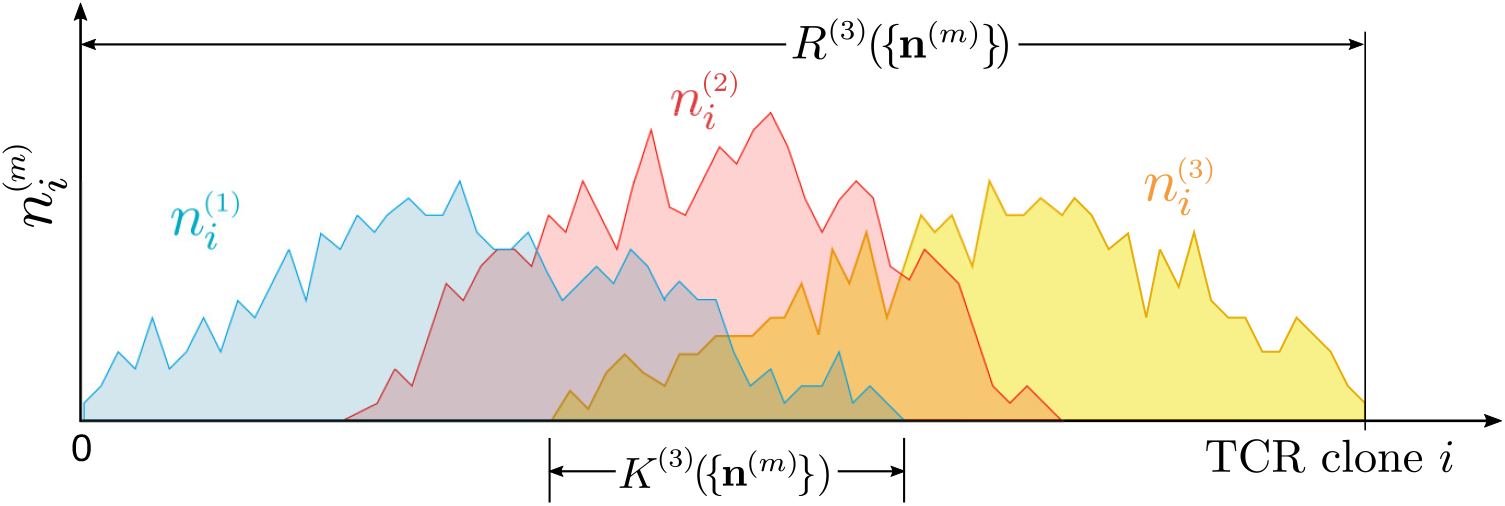
Three individuals with overlapping cell number distributions 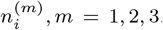. The richness R^(3)^ is the total number of distinct number of different TCRs found across all individuals, and is defined in Eq. (3.8). The overlap K^(3)^ is the number of TCR clones found in all three individuals, as defined in Eq. (3.17). For visual simplicity, the set of clones i present in each individual are drawn to be contiguous. When considering subsampling of cells from each individual, K^(M)^ will be reduced since some cell types i will be lost. The corresponding values, 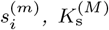, and 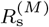 can be constructed from Eqs. (3.21) and (3.22) reflecting the losses from subsampling.

As with Eqs. (3.2) and (3.7), we can express the overlap in terms of the underlying individual probabilities **p**^(*m*)^ by weighting Eq. (3.17) by the *M*-population probability 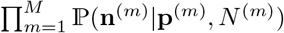 (see Eq. (3.1)) to find

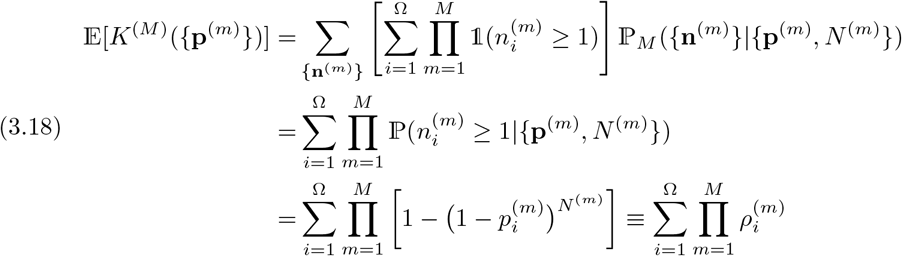

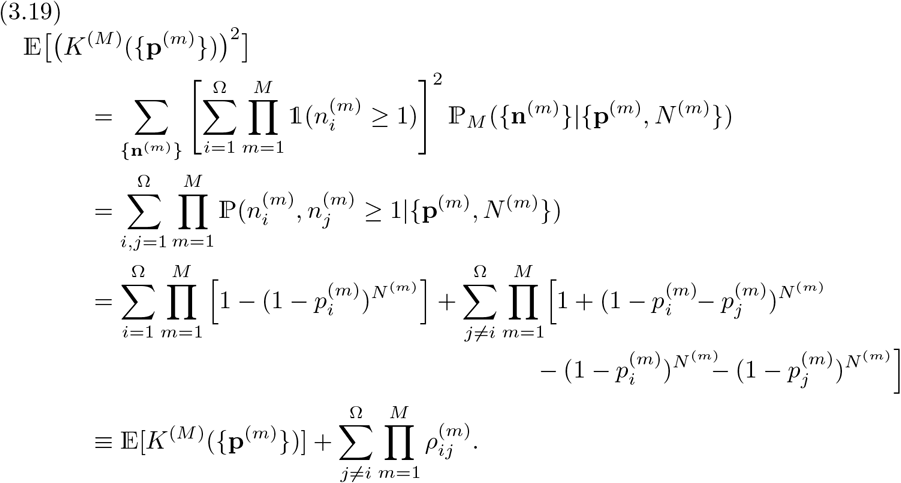

The expected number of clones shared among all *M* individuals, 𝔼[*K*^(*M*)^], provides a natural measure of *M*-publicness. Clearly, 𝔼[*K*^(*M*)^] *<* 𝔼[*K*^(*M*′)^] if *M > M*^′^. As with *M*-publicness, we can identify *M*-private clones as Ω − 𝔼 [*K*^(*M*)^], the expected number of clones that are not shared by all *M* individuals, *i*.*e*., that occur in at most *M* − 1 individuals. This “privateness” is related to a multi-distribution generalization of the “dissimilarity probability” of samples from two different discrete distributions [26]. Variations in *M*-publicness associated with a certain cell-type distribution are captured by the variance Var [*K*^(*M*)^] = 𝔼 [(*K*^(*M*)^)^2^] − 𝔼 [*K*^(*M*)^]^2^. If the total number of sequences Ω is very large, parallelization techniques (see Sec. 4) should be employed to evaluate the term 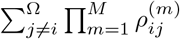 in 𝔼 [(*K*^(*M*)^)^2^].

A more specific definition of overlap or privateness may be that a clone must appear in at least some specified fraction of *M* tested individuals. To find the probability that *M*_*i*_ ≤ *M* individuals share at least one cell of a single type *i*, we use the Poisson binomial distribution describing independent Bernoulli trials on individuals with different success probabilities 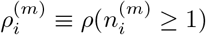:

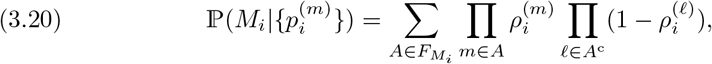

where 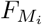 is the set of all subsets of *M*_*i*_ integers that can be selected from the set (1, 2, 3, …, *M*) and *A*^c^ is the complement of *A*. Equation (3.20) gives a probabilistic measure of the prevalence of TCR *i* across *M* individuals. For example, one can use it to define a mean frequency 𝔼[*M*_*i*_]*/M*. One can evaluate Eq. (3.20) recursively or using Fourier transforms, particularly for *M <* 20 [10, 27].

### 3.3. Subsampling

The results above are described in terms of the entire cell populations **n**^(*m*)^ or their intrinsic generation probabilities **p**^(*m*)^. In practice, one cannot measure 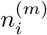 or even *N* ^(*m*)^ in any individual *m*. Rather, we can only sample a much smaller number of cells *S*^(*m*)^ ≪ *N* ^(*m*)^ from individual *m*, as shown in Fig. 2. Within this subsample from individual *m*, we can count the number 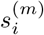 of type *i* cells. Since only subsamples are available, we wish to define quantities such as probability of occurrence, richness, and overlap in terms of the cell counts 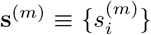 in the sample extracted from an individual. Quantities such as *sampled* richness and overlap can be defined in the same way except with **s**^(*m*)^ as the underlying population configuration. To start, first assume that the cell count **n** in a specific individual is given. If that individual has *N* cells of which *S* are sampled, the probability of observing the population **s** = {*s*_1_, *s*_2_, …, *s*_Ω_} in the sample is given by (assuming all cells are uniformly distributed and randomly subsampled at once, without replacement) [9]

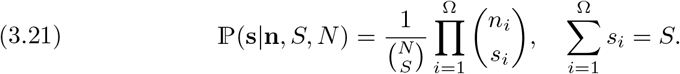

The probability that cell type *j* appears in the sample from an individual with population **n** can be found by marginalizing over all *s*_*j≠i*_, giving

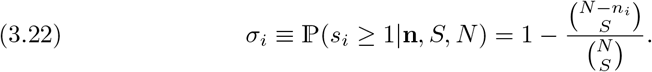

This result can be generalized to more than one TCR clone present. For example, the probability that both clones *i* and *j* are found in a sample is

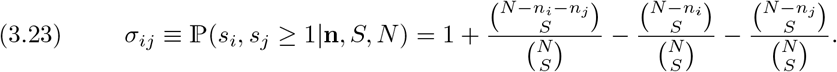

Using Eq. (3.21) as the probability distribution, we can also find the probability that clone *i* appears in any of the *M S*^(*m*)^-sized samples

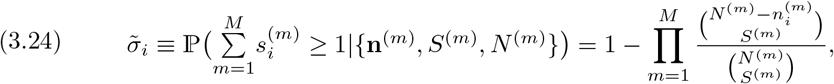

and the joint probabilities that clones *i* and *j* appear in any sample

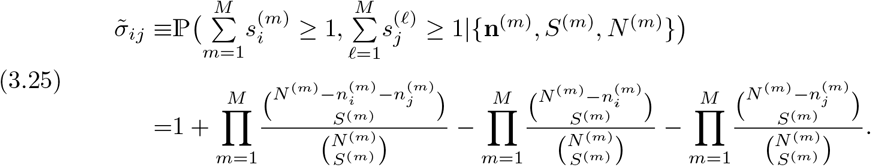

Quantities such as richness and publicness *measured within samples* from the group can be analogously defined in terms of clonal populations **s**^(*m*)^:

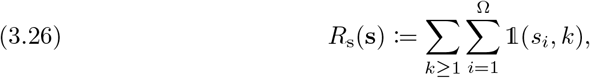

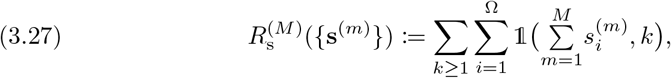

and

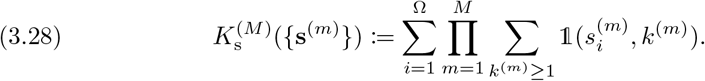

For a given **n**^(*m*)^, these quantities can be first averaged over the sampling distribution Eq. (3.21) to express them in terms of **n**^(*m*)^ and to explicitly reveal the effects of random sampling. The expected values of 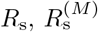, and 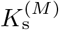 in terms of **n**^(*m*)^ can be found by weighting Eqs. (3.26), (3.27), and (3.28) by ℙ(**s**|**n**, *S, N*) and 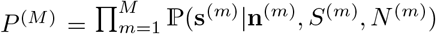:

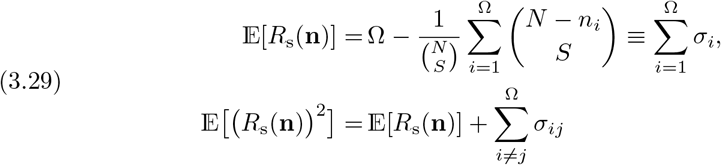

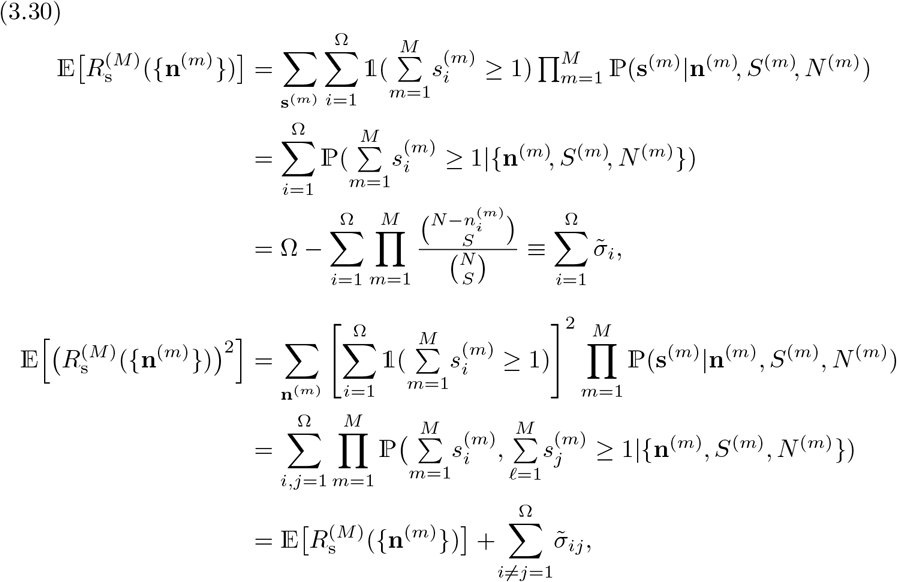

and

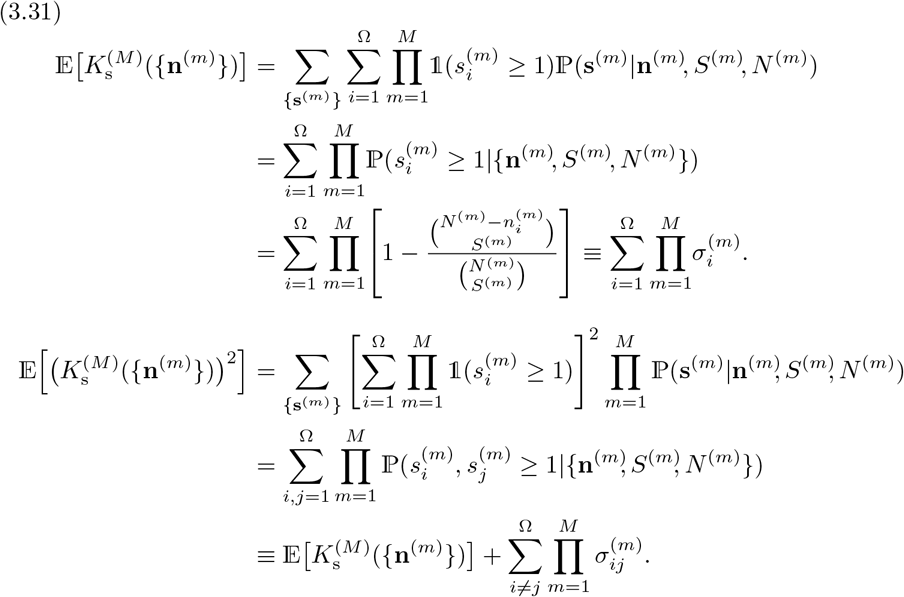

All of the above quantities can also be expressed in terms of the underlying probabilities **p**^(*m*)^ rather than the population configurations **n**^(*m*)^. To do so, we can further weight Eqs. (3.29), (3.30), and (3.31) over the probability Eq. (3.1) to render these quantities in terms of the underlying probabilities **p**^(*m*)^. However, we can also first convolve Eq. (3.21) with the multinomial distribution in Eq. (3.1) (suppressing the individual index *m*)

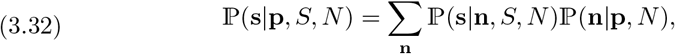

along with the implicit constraints 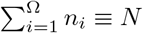 and 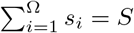 to find

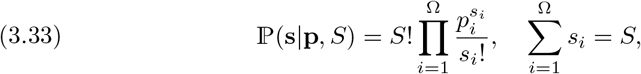

which is a multinomial distribution identical in form to ℙ(**n**|**p**, *N*) in Eq. (3.1), except with **n** replaced by **s** and *N* replaced by *S*. Uniform random sampling from a multinomial results in another multinomial. Thus, if we use the full multi-individual probability

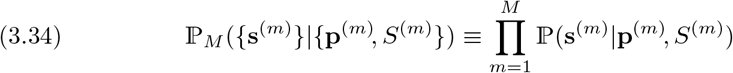

to compute moments of the sampled richness and publicness, they take on the same forms as the expressions associated with the whole-organism quantities. For example, in the **p** representation, the probability that clone *i* appears in the sample from individual *m* is

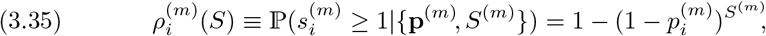

in analogy with Eq. (3.3), while the two-clone joint probability in the sampled from individual *m* becomes

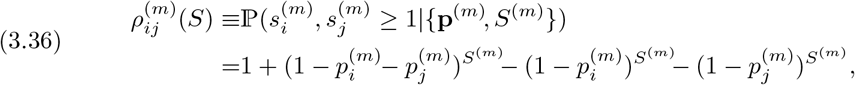

in analogy with Eq. (3.5). Similarly, for the overlap quantities, in analogy with Eqs. (3.10) and (3.13), we have

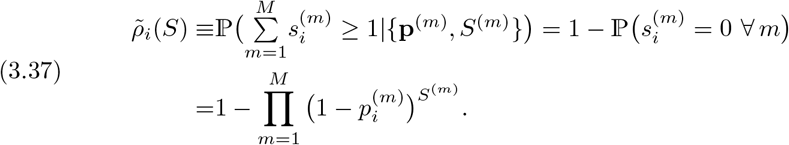

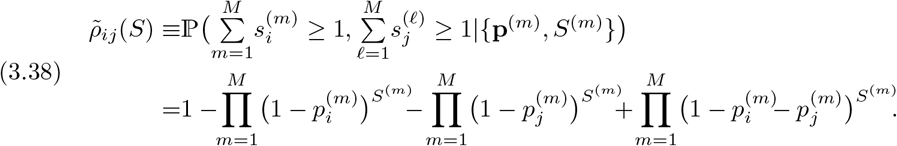

The expressions for the sampled moments 𝔼[*R*_s_(**p**)], 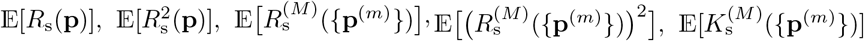, and 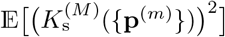 follow the same form as their unsampled counterparts given in Eqs. (3.7), (3.15), (3.16), (3.18), and (3.19), except with 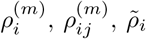, and 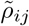 replaced by their 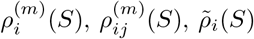, and 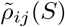 counterparts.

In addition to simple expressions for the moments of 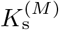, we can also find expressions for the probability distribution over the values of 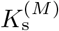. In terms of **n**^(*m*)^, since the probability that 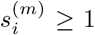 in the samples from all 1 ≤ *m* ≤ *M* individuals is 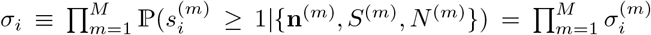, the probability that exactly *k* clones are shared by all *M* samples is

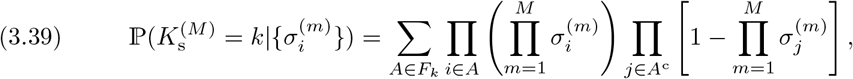

where *F*_*k*_ is the set of all subsets of *k* integers that can be selected from the set {1, 2, 3, …, *K*^(*M*)^} and *A*^c^ is the complement of *A*. Equation (3.39) is the Poisson binomial distribution, but this time the underlying success probabilities 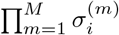 across all *M* individuals vary with TCR clone identity *i*.

Finally, inference of individual metrics from subsamples can be formulated. One can use the sampling likelihood function ℙ(**s**|**n**, *S, N*), Bayes rule, and the multinomial (conjugate) prior ℙ(**n|p**, *N*) to construct the posterior probability of **n** *given* a sampled configuration **s**:

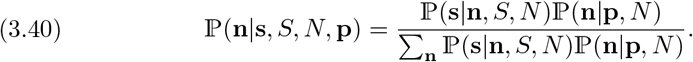

The normalization in Eq. (3.40) has already been found in Eqs. (3.32) and (3.33). Thus, we find the posterior

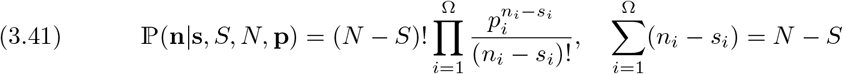

in terms of the hyperparameters **p**. Using this posterior, we can calculate the expectation of the whole organism richness 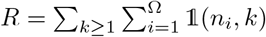,

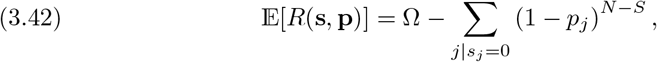

which depends on the sampled configuration only through the sample-absent clones *j*. Bayesian methods for estimating overlap between two populations from samples have also been recently explored [30].

## 4. Simulations

The sampling theory derived in the previous sections is useful for understanding the effect of different sampling distributions on measurable quantities such as the proportion of shared TCRs and BCRs among different individuals. Figures 4 and 5 show two examples of receptor distributions and corresponding relative overlaps for three individuals. First, to provide a simple, explicit example, we use three shifted uniform distributions in Fig. 4. In this example, the number of TCR or BCR sequences per individual is 100,000 and the group richness is *R*^(3)^ is 1,500. Based on the abundance curves shown in Fig. 4(a), we can readily obtain the maximum relative overlaps between individuals 1–3 (solid black, dashed blue, and dash-dotted red lines) and between all pairs of individuals. The maximum possible overlap between all three individuals and between individuals 1 and 3 is 500*/*1, 500 ≈ 0.33. For the two remaining pairs, the corresponding maximum overlap is 750*/*1, 500 = 0.5. In Fig. 4(b), we plot the normalized variance 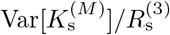 in the expected number of shared cells as a function of the number of sampled cells *S*^(*m*)^ from each individual. This variance reaches a maximum between 1,000 and 2,000 sampled cells, and decreases to negligible values when *S*^(*m*)^ ≳ 6, 000. Monitoring the variance can be used to monitor convergence of the measured number of shared cells. Using a number of sampled cells *S*^(*m*)^ from each individual that is smaller than the total number of cells per individual, we observe in Fig. 4(c) that the increase of 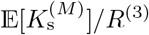 with *S*^(*m*)^ is well-described by Eq. (3.18).

**FIG. 4.**
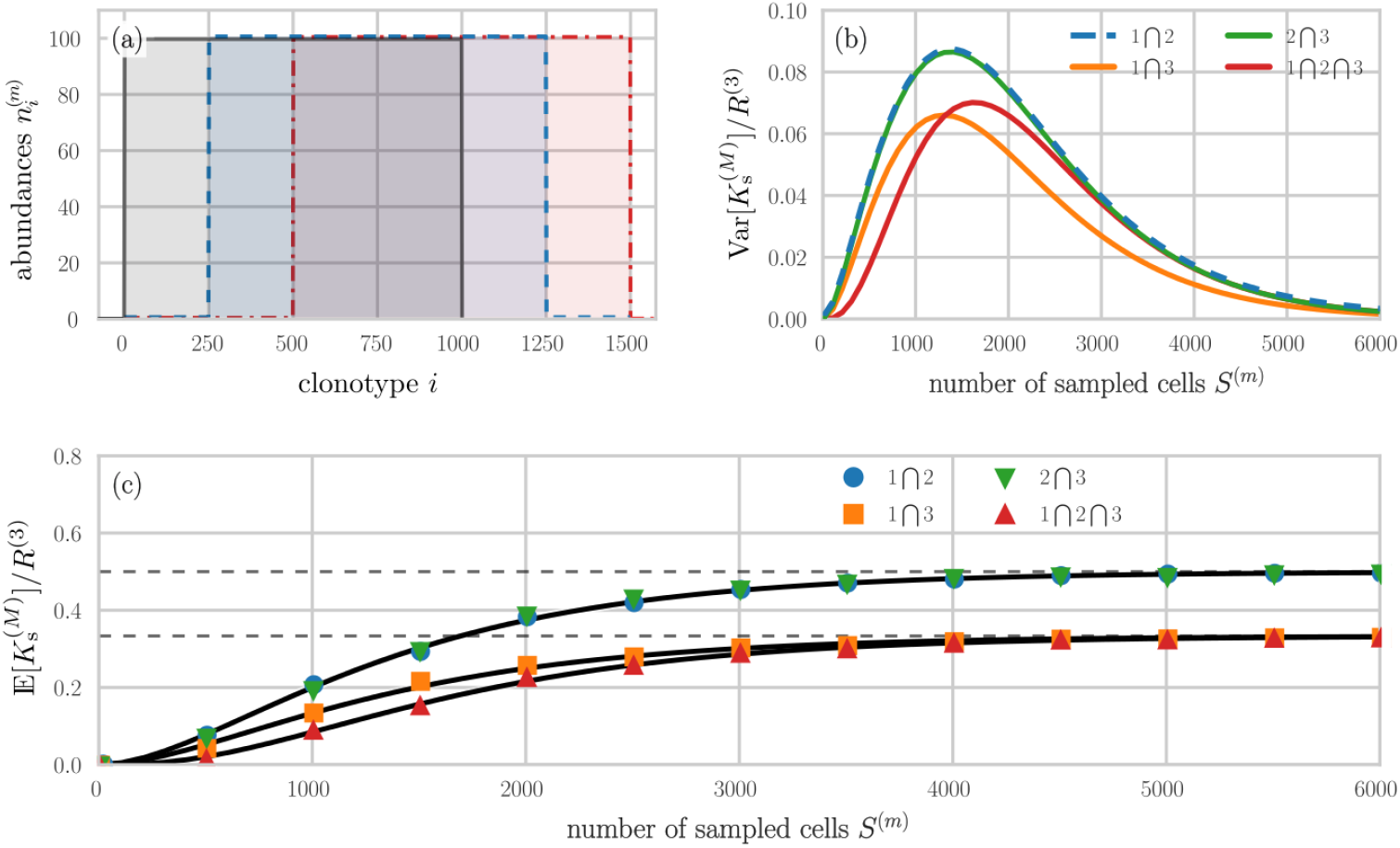
Sampling from shifted uniform distributions. (a) Synthetic TCR or BCR distributions of M = 3 individuals. The distributions in individuals 1, 2, and 3 are indicated by solid black, dashed blue, and dash-dotted red lines, respectively. Each individual has 100, 000 cells uniformly distributed across 1000 clones (100 cells per clone). The sampled group richness 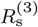 is 1, 500. (b) The variance 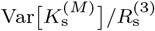 associated with the expected number of shared clones among M individuals. (c) Samples of size S^(m)^ have been generated to compute the relative overlaps between individuals 1 and 2 (blue disks), 2 and 3 (green inverted triangles), 1 and 3 (orange squares), and 1–3 (red triangles). The solid black lines in panel (c) show the corresponding analytical solutions 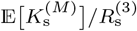 (see Eq. (3.18)). Dashed grey lines show the maximum possible relative overlaps 500/1, 500 ≈ 0.33 and 750/1, 500 = 0.5.

**FIG. 5.**
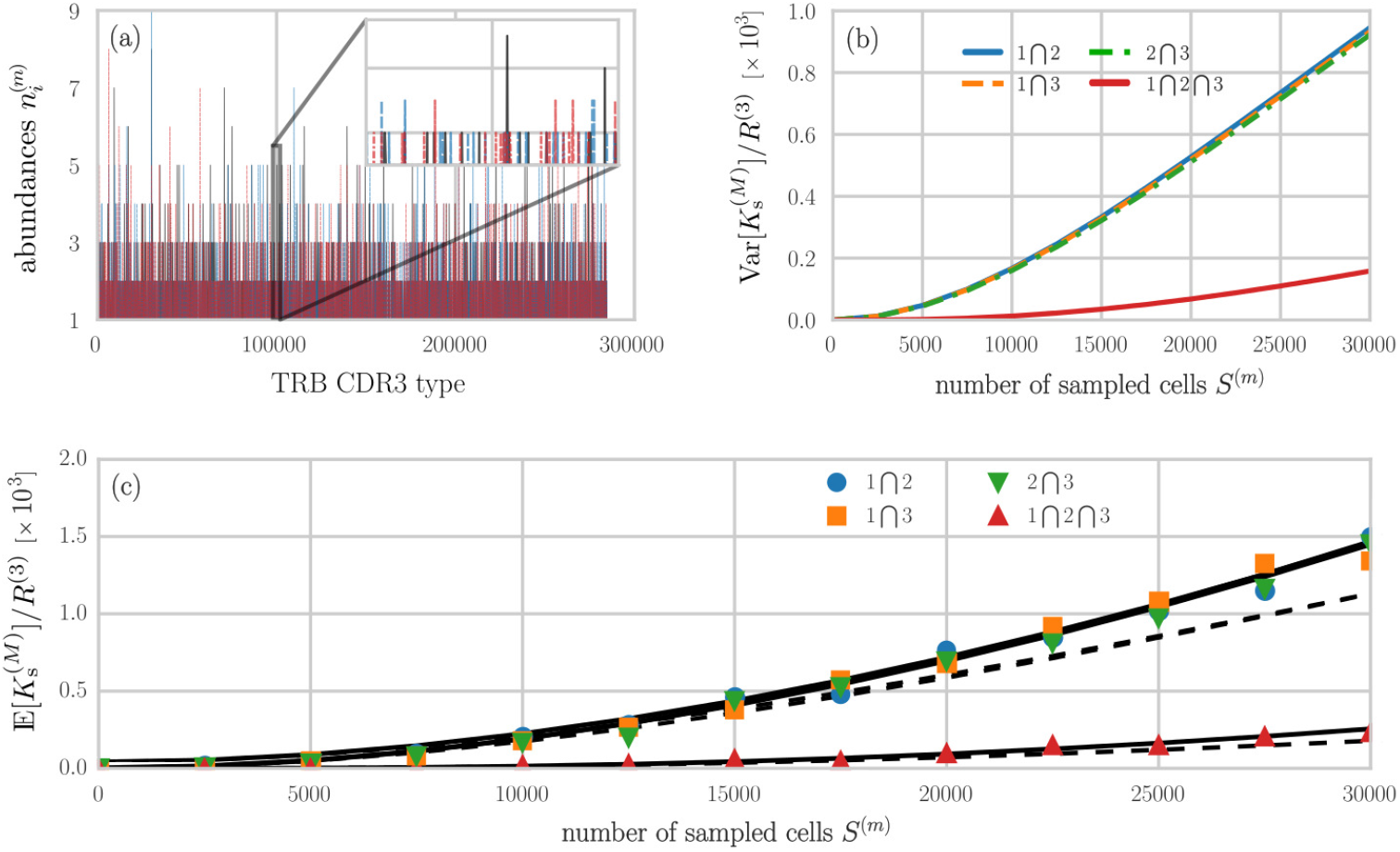
Sampling from empirical TRB CDR3 distributions and overlap metrics in the number representation. (a) Distributions of TRB CDR3 cells in M = 3 individuals. We used the SONIA package [17] to generate 100, 000 TRB CDR3 sequences for each individual. The total group richness R^(3)^ was found to be 284, 598. Equal sample sizes S^(m)^ were then drawn. (b) The variance 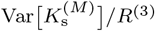 associated with the expected number of shared clones among M individuals, plotted as functions of S^(m)^. (c) Relative overlaps between individuals 1 and 2 (blue disks), 2 and 3 (green inverted triangles), 1 and 3 (orange squares), and 1–3 (red triangles). The solid black lines in panel (c) plot the corresponding analytical solutions 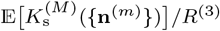 found in Eqs. (3.31). The dashed curve corresponds to using using the estimator 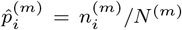 in the expression 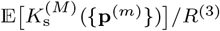 (Eq. (3.18) evaluated using 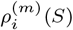 from Eq. (3.35)).

As an example of an application to empirical TRB CDR3 data, we used the SONIA package [17] to generate amino acid sequence data for three individuals with 10^5^ cells each. The combined richness across all individuals is *R*^(3)^ = 284, 598. We show the abundances of all sequences in Fig. 5(a). The corresponding variance associated with the measured number of shared cells and the expected number of shared cells are shown in Figs. 5(b,c), in the number representation. For a direct evaluation of Eq. (3.31), we evaluate the binomial terms in 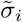 and 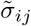 by expanding them according to, *e*.*g*.,

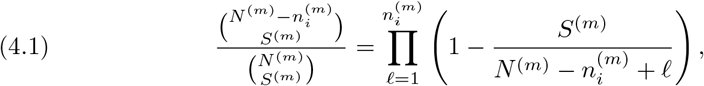

where *S*^(*m*)^*/N* ^(*m*)^ is the sample fraction drawn from the *m*^th^ individual. For large *n*_*i*_, other approximations, including variants of Stirling’s approximations can be employed for fast and accurate evaluation of binomial terms.

We can compare these number-representation results with the **p**-representation results by using the estimates 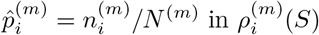 and 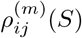 to compute the quantities in Eqs. (3.18) and (3.19). If the number of sampled cells *S*^(*m*)^ is not too large, the analytic approximation of using 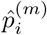 in 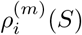 to calculate 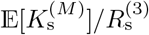 is fairly accurate, as shown by the dashed back curves in Fig. 5(c). Since the abundances of the majority of sequences are very small, finite-size effects lead to deviations from the naive approximation (3.35) (dashed black lines) as the numbers of sampled cells *S*^(*m*)^ grows large. Of course, we can also extract generation probabilities from SONIA and directly use Eq. (3.18) and 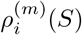 from Eq. (3.35) to find the **p**-representation *M*-overlap 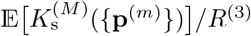.

Calculations were performed on an AMD^®^ Ryzen Threadripper 3970 using Numba to parallelize the calculation of Eqs. (3.31) and 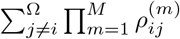 used in Var [*K*^(*M*)^].

## 5. Sampling resolution and information loss

We end with a brief discussion of information loss upon sampling and coarse-graining which arises when analyzing lower dimensional experimental/biochemical classifications of clones that are commonly used. Such lower-dimensional representations can be obtained through spectratyping [22, 19]. For TCRs, spectratyping groups sequences together and produces compressed receptor representations describing CDR3 length, frequency, and associated beta variable (TRBV) genes [21]. We quantify the information loss that is associated with (i) discretizing a continuous random variable, and (ii) coarse graining an already discretized distribution (*i*.*e*., spectratyping).

Given a discrete random variable *X* taking on possible values {*x*_1_, *x*_2_, …, *x*_Ω_}, let *p*_*i*_ = ℙ(*X* = *x*_*i*_). The entropy of this probability distribution is given by 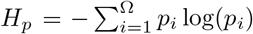. Similarly, one might define the differential entropy for a continuous random variable taking on values in the interval [*a, b*] as 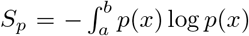. It is well-known that the differential entropy is not a suitable generalization of the entropy concept to continuous variables [28] since it is not invariant under change of variables and can be negative. These issues can be circumvented by introducing the limiting density of discrete points. Here, we present a more direct approach that will be sufficient for our application. For a probability density function 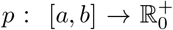 we introduce a discretizing morphism 𝒟_Δ_ so that

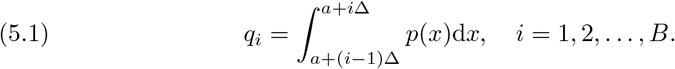

describes a random variable taking on values in each of the (*b* − *a*)*/*Δ = *B* bins.

To quantify the amount of information lost in this discretization step, consider the entropy 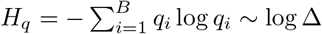 in the Δ → 0 limit. If we want to evaluate any information loss as a difference between the (finite) differential entropy *S*_*p*_ and the (diverging) entropy *H*_*q*_ we need to account for this logarithmic contribution by defining the corresponding information loss as

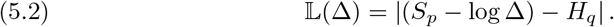

By absorbing the logarithmic contribution into the differential entropy, we find the correct continuous entropy according to Jaynes [28] using the limiting density of discrete points.

As an example, we compute the information loss associated with discretizing the truncated power law

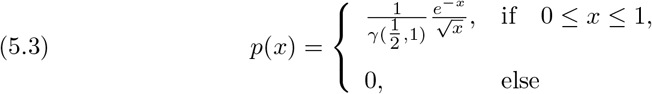

where 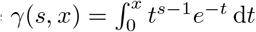 is the lower incomplete gamma function. The distribution Eq. (5.3) gives rise to few high-abundance clones and many low-abundance clones, as typical for TCR receptor sequences [44]. Analytic expressions for the discretized probabilities *q*_*i*_ are lengthy so we numerically compute *q*_*i*_ to evaluate the information loss 𝕃(Δ). Eq. (5.2) is plotted as a function of the number of bins *B* = 1*/*Δ in Fig. 6(a). The information loss decreases with the number of bins, as this results in the discrete distribution gathering more information about its continuous counterpart.

**FIG. 6.**
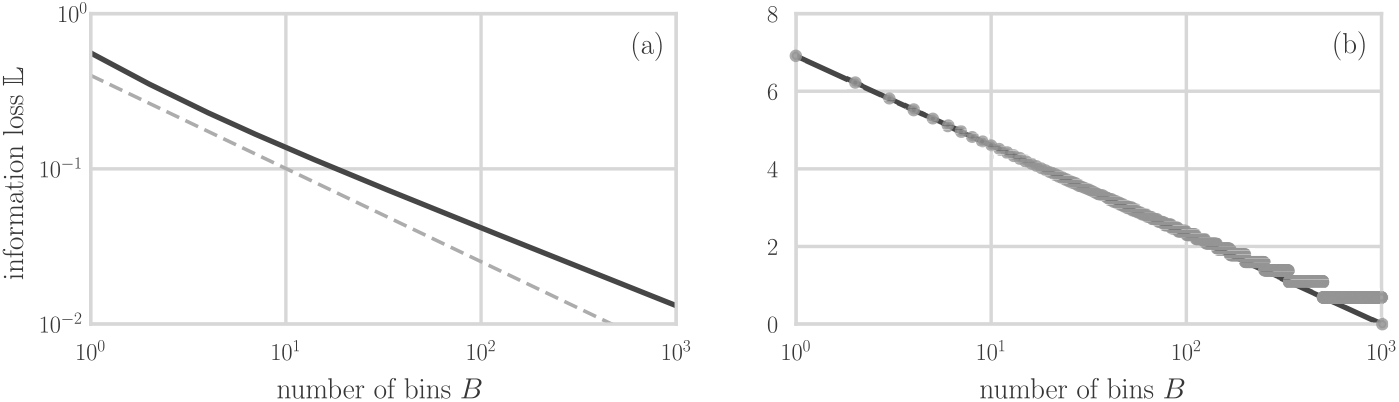
The information loss 𝕃 as a function of the number of discretization bins B. The loss is least as the number of integration bins B → ∞. (a) The solid black line shows the information loss associated with discretizing a truncated power law [see Eq. (5.3)], and the dashed grey line is a guide-to-the-eye (power law) with slope − 0.6. (b) Grey disks show the information loss associated with coarse graining a discrete and uniform random variable with initially Ω = 1, 000 traits. The solid curve shows the corresponding analytical result for the difference in information 𝕃 = − log(B/Ω) between discretizing a continuous uniform distribution by Ω and by B bins.

While the connection between continuous probability distributions and their discretized counterparts has important consequences for sampling, information loss also occurs in spectratyping when an already discrete random variable is coarse grained. In this scenario, the information loss can be quantified uniquely (up to a global multiplicative constant) by the entropy difference of the distributions [4]. The difference between the full 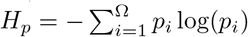 and the coarse-grained 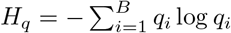 can be explicitly evaluated for uniformly distributed probabilities.

For any number *B <* Ω we can define a coarse graining procedure that yields only *B* traits by defining the bin size Δ = ceil(Ω*/B*) and grouping together Δ traits into each bin. The last bin might be smaller than the other bins or even empty. The information loss of this procedure is shown in Fig. 6(b) for an initially uniform distribution of Ω = 1, 000 traits. Across certain ranges of *B*, plateaus can build since our coarse graining might add zero probabilities. However, we can instead start from a continuous distribution and compare the discretization with Ω = 1, 000 bins to any other binning with *B* ≤ Ω.

Comparing a coarse-grained uniform distribution with *B* bins to the discretized distribution with Ω bins yields the information loss with respect to the initial discrete distribution 𝕃 = − log(*B/*Ω) ≥ 0. We plot this analytical prediction against the information loss L associated with coarse graining an already discrete distribution in Fig. 6(b), showing them to be well-aligned.

## 6. Discussion and Conclusions

Quantifying properties of cell-type or sequence distributions is an important aspect of analyzing the immune repertoire in humans and animals. Different methods have been developed to estimate TCR and BCR diversity indices such as the total number of distinct sequences in an organism (*i*.*e*., species richness) [29, 44]. Another quantity of interest is the number of clones that are considered “public” or “private,” indicating how often certain TCR or BCR sequences occur across different individuals.

Public TCR*β* and BCR sequences have been reported in a number of clinical studies [34, 35, 38, 40, 5, 39]. However, the terms “public” and “private” clonotypes are often based on different and ambiguous definitions. According to [38], a “public sequence” is a sequence that is “*often* shared between individuals” [38], while [25] refers to a sequence as public if it is “shared across individuals”. In addition to ambiguities in the definition of what constitutes a private/public sequence, overlaps between the immune repertoires of different individuals are often reported without specifying confidence intervals, even though variations may be large given small sample sizes and heavy tailed sequence distributions.

In this work, we provided mathematical definitions for “public” and “private” clones in terms of the probabilities of observing a number of clones that are observed in all *M* randomly chosen individuals. Besides defining individual repertoire probability distributions, our results include analytic expressions for individual and multi-individual expected richness and expected overlap as given by Eqs. (3.2), (3.7), (3.15), (3.17), and (3.18). The corresponding predictions for the expected richness and expected overlap in subsamples were also found, and are summarized in Eqs. (3.26), (3.27), (3.28), (3.29), (3.30), and (3.31). The variability of quantities (second moments) such as the *M*-overlap and subsampled overlap were also derived. Our results are summarized in Table 1 where we provide expectations and second moments of all quantities as a function the cell population configurations **n**^(*m*)^ or as a function of the underlying clone generation probabilities **p**^(*m*)^, as is generated by models such as SONIA [17]. These quantities can also be straightforwardly computed in the “clone count” **c**-representation, which we leave as an exercise for the reader.

Further inference of richness and overlap given sample configurations can be developed using our results. For example, the parametric inference of expected richness in an individual given a sampled configuration **s** can be found using the multinomial model and Bayes rule, as presented in Eq. (3.42). All of our results assume knowledge of at least one parameter Ω. However, this global intrinsic richness often also needs to be estimated or modeled. Numerous parametric and nonparametric approaches have been developed in the statistical ecology literature [8, 43, 23, 13, 24, 11, 9, 7], as well as expectation maximization methods to self-consistently estimate richness and most likely clone population **n** [29]. Finally, in the context of coarse-graining, or spectratyping [12], we have discussed methods that are useful to quantify the information loss associated with different levels of coarse graining TCR and BCR sequences.

